# *Pseudomonas aeruginosa* PA5oct jumbo phage impacts planktonic and biofilm population and reduces its host virulence

**DOI:** 10.1101/405027

**Authors:** Tomasz Olszak, Katarzyna Danis-Wlodarczyk, Michal Arabski, Grzegorz Gula, Barbara Maciejewska, Slawomir Wasik, Cédric Lood, Gerard Higgins, Brian J. Harvey, Rob Lavigne, Zuzanna Drulis-Kawa

## Abstract

In this work we analyzed the impact of jumbo phage PA5oct on the planktonic, cell line adhered, and biofilm population of *P. aeruginosa*. PA5oct has a broad host-range, able to infect up to 40% of our clinical *P. aeruginosa* Cystic Fibrosis (CF) collection. In the airway surface liquid (ASL) model, the infection of PA5oct effectively reduced the bacterial population both adhered to epithelial cells, mucus entrapped, and dispersed. The explanation for its infectivity can also be linked to the sensitization of infected bacteria to the innate immune mechanisms and pro-inflammatory effect. Interferometry of a 72-hour old biofilm highlighted the contribution of PA5oct in biofilm matrix degradation. Interestingly, two virion-associated proteins, gp162 and gp205, have been found as putative enzymes that can degrade matrix exopolysaccharides. Two third of biofilm clones developed PA5oct phage-resistance and the cross-resistance to both LPS- and pili-dependent phages. Simultaneously, all clones resistant to phage PA5oct maintain the phage DNA within the population, strongly reducing bacterial virulence *in vivo*. These properties can be considered as key parameters for the application of this bacterial virus in phage therapy settings.

**Originality-Significance Statement:** The emergence of phage-resistant mutants is a key aspect of lytic phages-bacteria interaction and the main driver for the co-evolution between both organisms. However, this fundamental property also has implications for bacterial eradication in phage therapy settings. Here, we analyze the impact of PA5oct jumbo phage treatment of planktonic/cell line associated and sessile *P. aeruginosa* population in a preclinical evaluation of this phage for therapeutic applications. Besides its broad-spectrum activity and efficient bacteria reduction in both airway surface liquid (ASL) model, and biofilm matrix degradation, PA5oct appears to persist in most of phage-resistant clones. Indeed, a high percentage of resistance (20/30 clones) to PA5oct is accompanied by the presence of phage DNA within bacterial culture. Moreover, the maintenance of this phage in the bacterial population is correlated to reduced *P. aeruginosa* virulence, coupled with a sensitization to innate immune mechanisms, and a significantly reduced growth rate. We observed rather unusual consequences of PA5oct infection causing an increased inflammatory response of monocytes to *P. aeruginosa*. This, phenomenon combined with the loss or modification of the phage receptor makes most of the phage-resistant clones significantly less pathogenic in *in vivo* model. During phage therapy treatment, phage-resistance is considered as an adverse effect, but our results indicate that it leads to diminished bacterial virulence and increased clearance of the infected host. These findings provide new insights into the general knowledge of giant phages biology and the impact of their application in phage therapy.

## Introduction

Finding a solution to the antibiotic resistance problem is one of the greatest challenges of modern science and medicine, and the search for alternative strategies to antibacterial therapy has led to a renewed appreciation of bacteriophages (Wittebole *et al.*, 2014). Phage therapy efficacy studies (Galtier *et al.*, 2017; Roach *et al.*, 2017; Forti *et al.*, 2018) and recent advances in the regulatory frame, in which phage therapy can be adopted as part of ‘Magistral preparations’ (Pirnay *et al.*, 2018) have shifted the focus from proving the efficacy of phage therapy to its operational implementation, while expanding the number of phage isolates available the design of therapeutic cocktails.

Among the phages currently evaluated for therapeutic applications are the jumbo phages, defined by long dsDNA genomes in excess of 200kb (Hendrix, 2009). Jumbo phages are commonly found in (Krylov *et al.*, 2012) commercial phage therapy products, owing to their broad host range. The large coding potential of jumbo phage allows them to be (partly) independent from the bacterial enzymes, empowering their expanded host range (Yuan and Gao, 2017). However, some Jumbo phages are marked with a high frequency of transduction, which impacts the evolution of bacteria and raises questions on the safety of their use in therapy (Monson *et al.*, 2011). The first jumbo phage was discovered over 40 years ago (*Bacillus* bacteriophage G), but the frequency of giant phages isolation remains rather low (less than 90 complete genomes in GenBank database). They form an incredibly diverse group, and new isolates of jumbo phages generally show low similarity to those present in public databases. They are also poorly characterized functionally, with annotated genomes that contain a vast majority of genes with undefined function (Lecoutere *et al.*, 2009).

The first sequenced jumbo phage genome specific for *Pseudomonas aeruginosa* was phiKZ - a giant lytic myovirus with a broad host range, isolated in Kazakhstan. Its large capsid (120 nm in diameter) encloses a linear, circularly permuted, terminally redundant genome (280,334 bp, 36.8% G+C) and is capable of carrying large fragments of the bacterial DNA (generalized transduction) (Mesyanzhinov *et al.*, 2002). This phage has become a hallmark example for structural analysis of phage particle and for genetic and structural analysis (Fokine *et al.*, 2007; Ceyssens *et al.*, 2014; Chaikeeratisak *et al.*, 2017). Currently, over twenty giant *Pseudomonas* bacteriophages within the diverse *Phikzvirus* genus have been isolated(Krylov *et al.*, 2007; Monson *et al.*, 2011; Danis-Wlodarczyk *et al.*, 2016).

*P. aeruginosa* phage PA5oct was isolated from sewage samples in Wroclaw, Poland. It is a representative of completely new genus of the *Myoviridae* family. The analysis of virion morphology (TEM micrograph) and genome size ranks PA5oct phage among the largest known bacterial viruses. The head diameter of PA5oct is about 131 nm and its tail is about 136 nm long (Drulis-Kawa *et al.*, 2014). It has a linear dsDNA genome containing 286,783 bp, making it the third largest genome of *Pseudomonas* phage (Genbank MK797984), and for which a comprehensive temporal transcriptome analysis, structural proteomics analysis and host transcription response has been studied (Integrative omics analysis of *Pseudomonas aeruginosa* virus PA5oct highlights the molecular complexity of jumbo phages. 2019. Lood C, Danis-Wlodarczyk K, Blasdel B, Jang HB, Vandenheuvel D, Briers Y, Noben JP, van Noort V, Drulis-Kawa Z, Lavigne R, bioRxiv 679506; doi: 10.1101/679506).

In this study we assess the influence of PA5oct on a population of *Pseudomonas aeruginosa*, both planktonic and sessile. We apply an integrated approach for the *in vitro* preclinical evaluation of its therapeutic potential using an established laser interferometry technique to measure the biofilm matrix degradation, and an advanced Airway Surface Liquid infection model that mimics *in vitro* the normal and CF lung environments. We also analyze the short- and long-term consequences of an apparent process of phage maintenance in the bacterial population, focusing on the emergence of phage-resistant clones, cross-resistance to other non-related phages and the influence of PA5oct on bacterial virulence.

## Results

### The host range of phage PA5oct suggests an increased activity against clinical CF isolates

The lytic activity of PA5oct was examined on two independent *P. aeruginosa* panels. First, on the COST international reference panel of 43 clinical *P. aeruginosa* (Cullen *et al.*, 2015). On that collection, phage PA5oct infects 24% of the isolates, a rate that is more limited when compared to representatives of the *Luz7virus* LUZ7 (42%), *Phikmvvirus* group (LUZ19 44%), *Pbunavirus* group (LBL3 40%, KT28 28%, KTN6 42%) and *Phikzvirus* group (phiKZ 47%, KTN4 33%). A second collection of 47 cystic fibrosis isolates obtained from the Leuven University hospital, Leuven, Belgium shows different results (Table S1-S2). The activity on these isolates from long term chronic infections was much broader, with PA5oct showing productive infection on 40% of the collection, whereas phages such as phiKZ and LUZ19 have a more narrow host-range (20% and 21% respectively). The COST international reference panel also contains 25 CF isolates but PA5oct phage only lyses six of them. No correlation could be observed in terms of phage activity versus CF early/late type of *P. aeruginosa* isolate. These results indicate the antibacterial potential of PA5oct against clinical *P. aeruginosa* strains isolated from different kinds of infections (burn wounds, CF-patients pneumonia, nosocomial pneumonia, and urinary tract infection).

Propagation experiments on a panel of specific PAO1 cell wall knock-outs reveals that PA5oct requires the presence of LPS and at least a second host cell surface receptor, like the Type IV pili (Table 1). The flagella mutant ΔfliC wt algC wt pilA does not provide conclusive results concerning the susceptibility to phage infection.

**Table 1.**
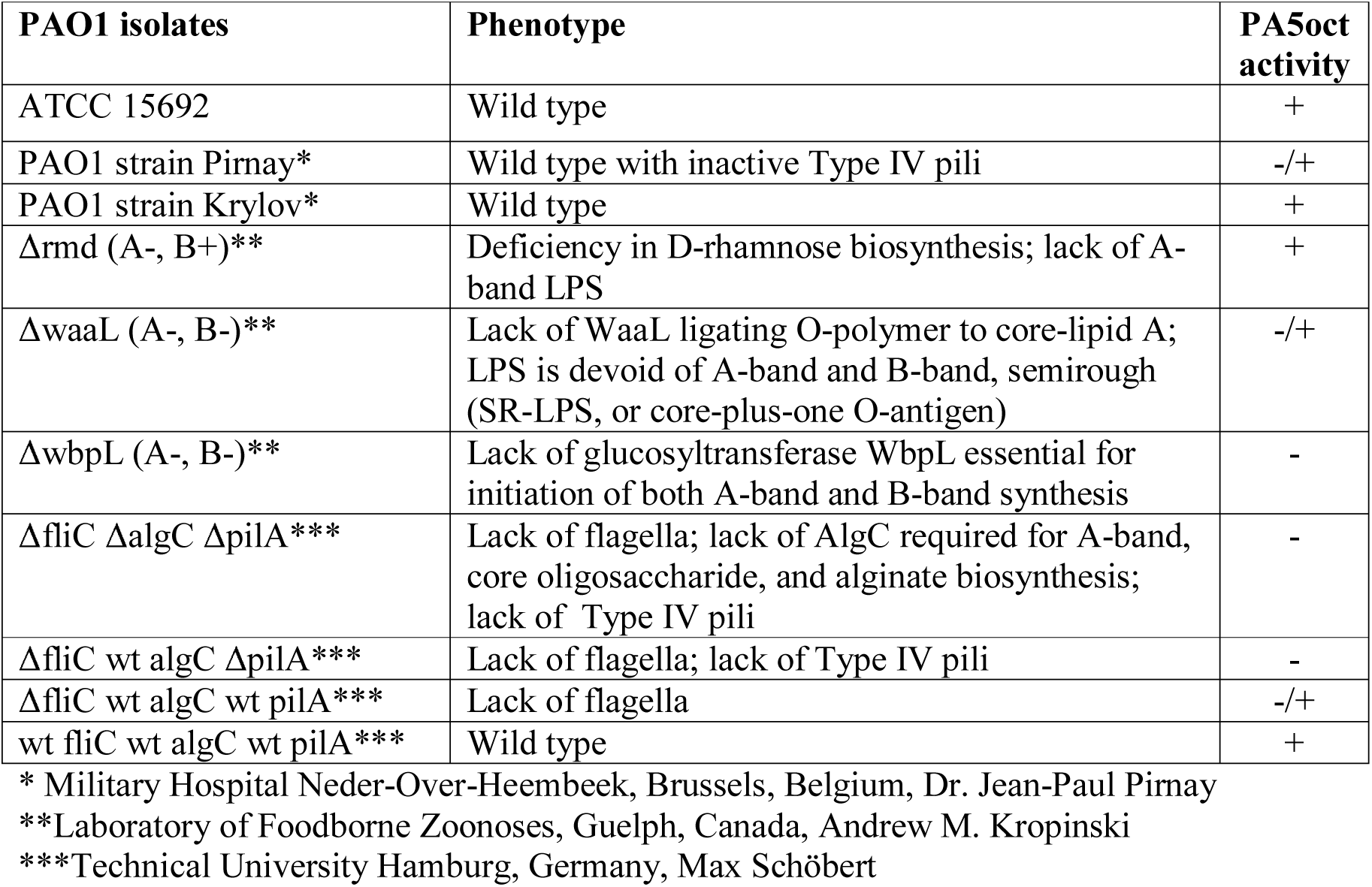
Phage receptor identification on *P. aeruginosa* PAO1 mutants.

### Phage PA5oct infection causes significant reduction of planktonic/cell line-associated bacteria in an Airway Surface Liquid infection model

The *in vitro* antibacterial activity of PA5oct against *P. aeruginosa* was assessed using the ASL model on normal NuLi-1 and cystic fibrosis CuFi-1 bronchial epithelium cell lines (Zabner *et al.*, 2003; Danis-Wlodarczyk *et al.*, 2016). Both cell lines mimic the natural environment of *Pseudomonas* lung infection both in healthy and cystic fibrosis patients respectively. Three different, well-characterized *P. aeruginosa* strains were applied, including the model PAO1, a burn-infection strain nonCF0038, and a small colony variant CF708 that was isolated from the late stage of CF infection (Olszak *et al.*, 2015). Epithelial cells viability controls were established as well, and no toxicity influence was observed for phage and bacterial samples.

After infection of both epithelial cell lines for 3 h, the colony count showed that all *P. aeruginosa* strains efficiently propagated in both ASL setups (10^7^ - 10^9^ cfu/ml). Phage treatment significantly (p < 0.05) reduced the CFU counts for normal NuLi-1, where 4.5 log, 6.5 log and 3 log decreases were observed for PAO1, nonCF0038 and CF708, respectively (Fig. 1A). The phage application for CuFi-1 epithelia infection was also very effective (p < 0.05) giving 5 log, 2.5 log and 5 log reductions in CFU of PAO1, nonCF0038 and CF708, respectively (Fig. 1B).

**Figure 1.**
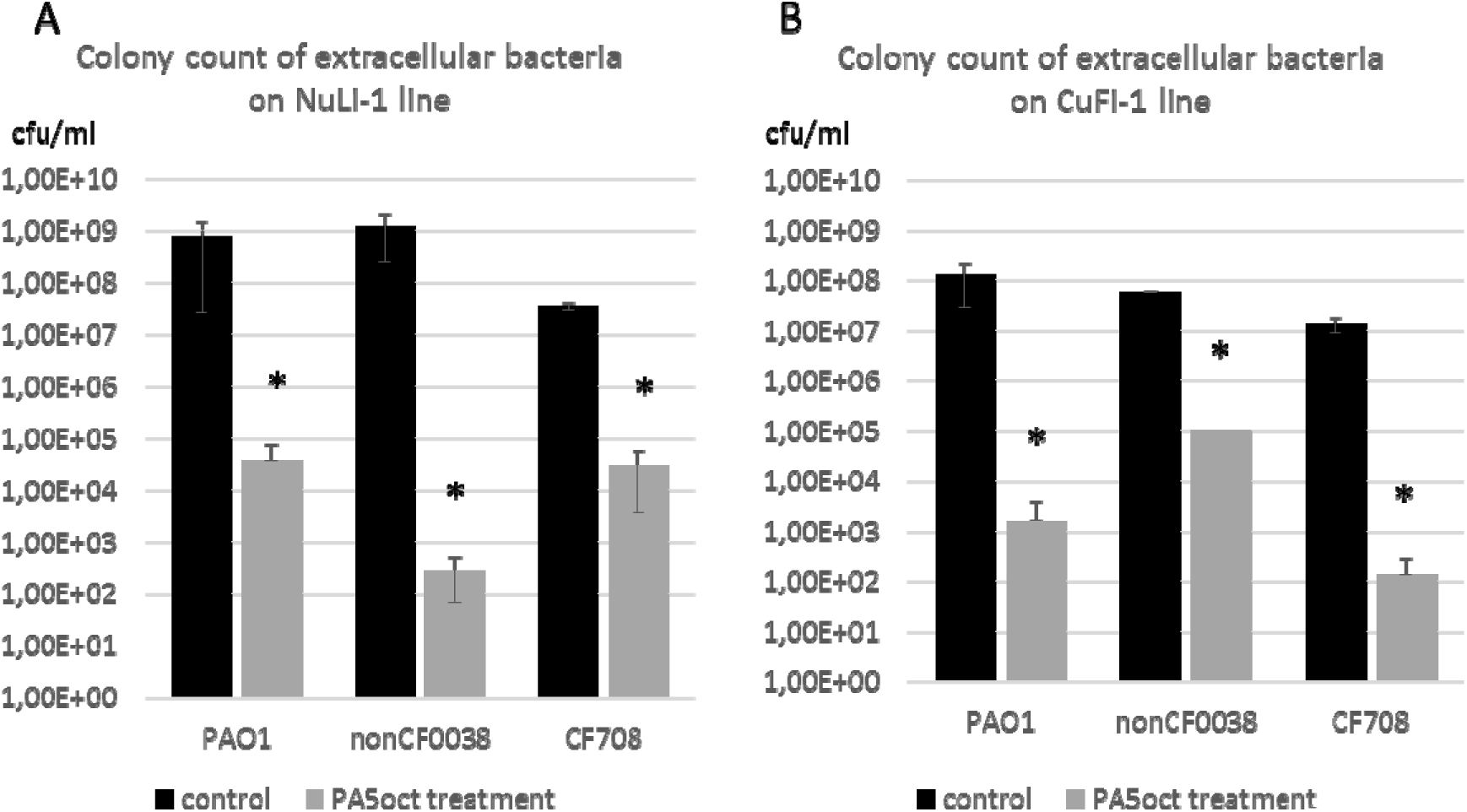
Phage PA5oct treatment of *P. aeruginosa* infecting NuLi-1 (A) and CuFi-1 epithelial cells (B). Colony count of bacteria collected from apical wash. The gray bars represent bacteria titers after 1.5 h of incubation with PA5oct phage. The black bars represent controls without phage treatment. The error bars indicate the standard deviation. The results are presented as the means ± SD. Statistical analysis was made using the ANOVA test (denoted *p*-values < 0.05). (*) *p*-values < 0.05.

### Real time measurement of P. aeruginosa biofilm diffusion properties

The direct effect of PA5oct phage on the mature (72h) biofilm *P. aeruginosa* PAO1 (overspread on hydrophilic Nephrophane membrane scaffolding) was investigated using laser interferometry, a standardized approach used for diffusion properties measurement, which allows better comparison to other experiments on how the biofilm biomass is degraded (Danis-Wlodarczyk *et al.*, 2015, 2016). First, the membrane was examined in terms of the biofilm coverage. Photos of membranes stained with crystal violet (CV) were collected and converted into grey-scale digital images (1 denotes black and 256 denotes white) (Fig 2A). Using the ImageJ computer imaging software (Schneider *et al.*, 2012) the degree of membrane coverage by biofilm was estimated at around 93%. Second, the biofilm was treated with active or inactive phages for 4 hours and then washed out to remove phage suspensions. Next, the diffusion event was measured by interferometry for 40 min in real-time (Fig. 2B). An increase of TSB diffusion rate through the biofilm layer correlates with the structural degradation of the biofilm/matrix. The diffusion rate of medium transported through the intact biofilm-covered membrane after 40 min (0.605 mg) was significantly lower than for biofilm (p < 0.05) after active and inactivated phage treatment, reaching 1.64 mg and 1.17 mg, respectively. This experiments indicated that phage PA5oct as infecting virions as well as protein particles was able to reduce the density of *Pseudomonas* biofilm after 4 h-treatment. The increase in diffusion obtained after the application of inactivated particles may be explained by an enzymatic activity of virion-associated proteins responsible for biofilm matrix degradation. Although, the plaques produced by PA5oct phage are very small with no halo zones, the *in silico* analysis of PA5oct virion-associated proteins using Phyre2, UniProt BLAST, BLASTP, HMMER, and SwissModel (EsPASy) software allowed selecting of potential phage enzymes able to degrade bacterial exopolysaccharide matrix. The gp162 was the first hit with 100.0% confidence in the model (N-term: tail connector protein/centre: transferase/C-term: lyase (351-451aa) forming putative tail sheath stabilizer or envelope glycoprotein. The second hit was gp205 a putative hydrolase alpha-n-acetylglucosaminidase (66.5% confidence in the model) catalyzing the hydrolysis of glycosidic bonds in complex sugars. In the case of halo-exhibiting phages, mostly *Podoviridae*, the zone occurs as an effect of depolymerase diffusion or phage diffusion in the agar. In the case of giant phages with virion-associated enzymes the diffusion in the agar is strongly limited by the size of phage capsid, thus the halo zone might not be observed.

**Figure 2.**
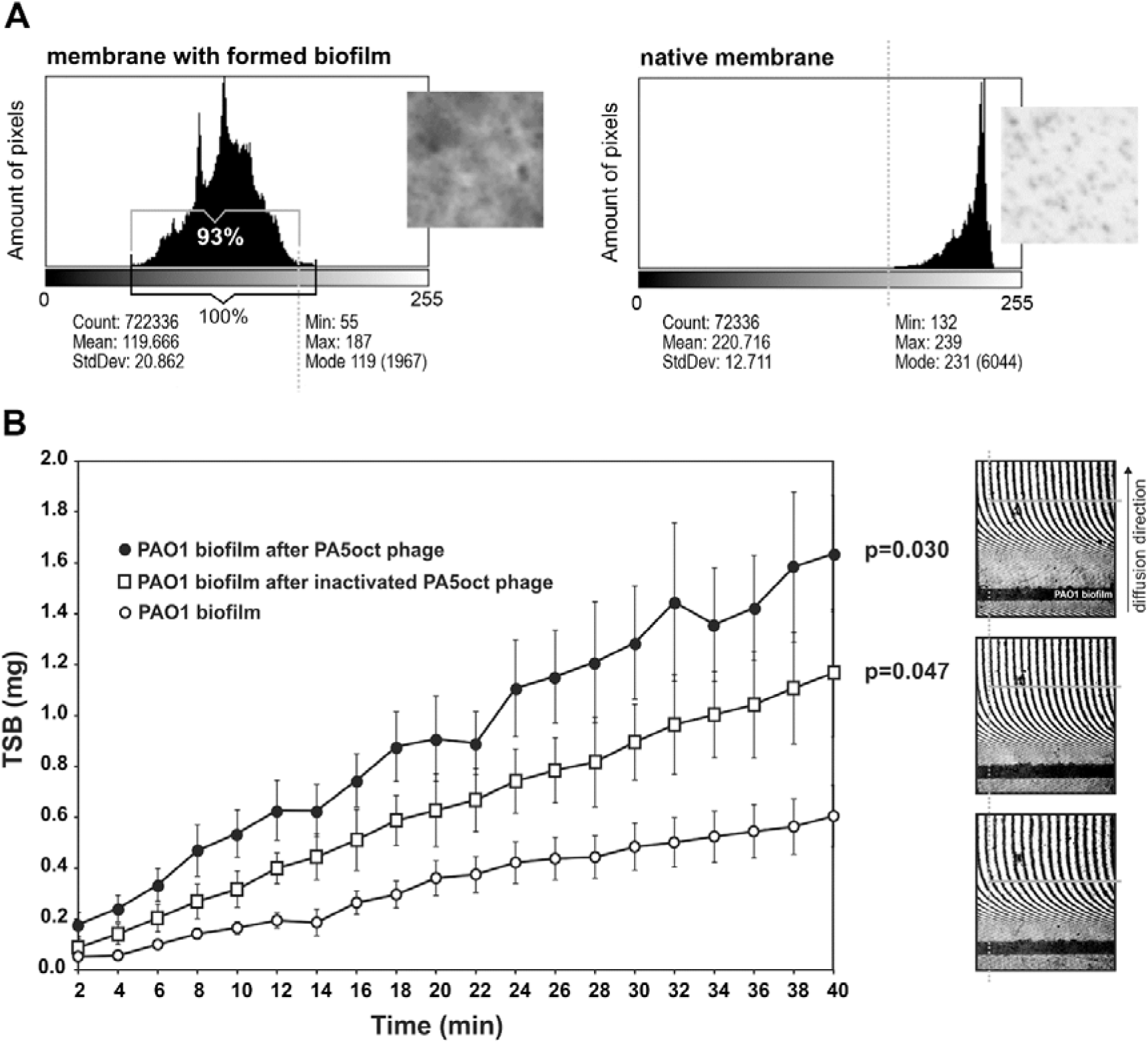
Real time measurement of biofilm permeability, after 4 h of phage treatment. (A) Membrane coverage analysis by CV staining and ImageJ imaging software. Native membrane was used as a control. (B) Laser interferometry analysis of TSB medium diffusion through PAO1 biofilm treated with PA5oct phage. Untreated biofilm was used as a control. Error bars denote SD. The results displayed are the mean of three independent experiments. Statistical analysis was made by the ANOVA test to compare data of treated biofilm versus control native biofilm at 40 min time point. The examples of interferograms (40 min.) for PAO1 biofilm treated with active and inactivated PA5oct phages as well as control (from the top of a right-hand panel).

To establish the antimicrobial activity of phage PA5oct against sessile bacteria, three complementary assays have been performed: the biomass CV staining and the measurement of pyocyanin and pyoverdin/pyochelin secretion (Fig. 3). Experiments were performed on a Nephrophane membrane with overgrown PAO1 biofilm, at various time points (24, 48 and 72 h). The CV staining of biofilm biomass showed a significant effect of the active PA5oct against mature biofilm (72 h). Moreover, the analysis of pyocyanin and pyoverdin/pyochelin secretion indicated that active phages significantly decreased the level of these compounds in the biofilms tested (72 h), whereas UV-inactivated phages had no significant effect. A positive correlation between biofilm formation, pyocyanin and pyoverdin/pyochelin levels was observed in the supernatant, indicating that a reduced level of *Pseudomonas*-specific compounds was related to phage activity. In summary, we show that phage PA5oct primarily affects mature biofilm, reducing its biomass as well as inhibits the production of selected virulence factors.

**Figure 3.**
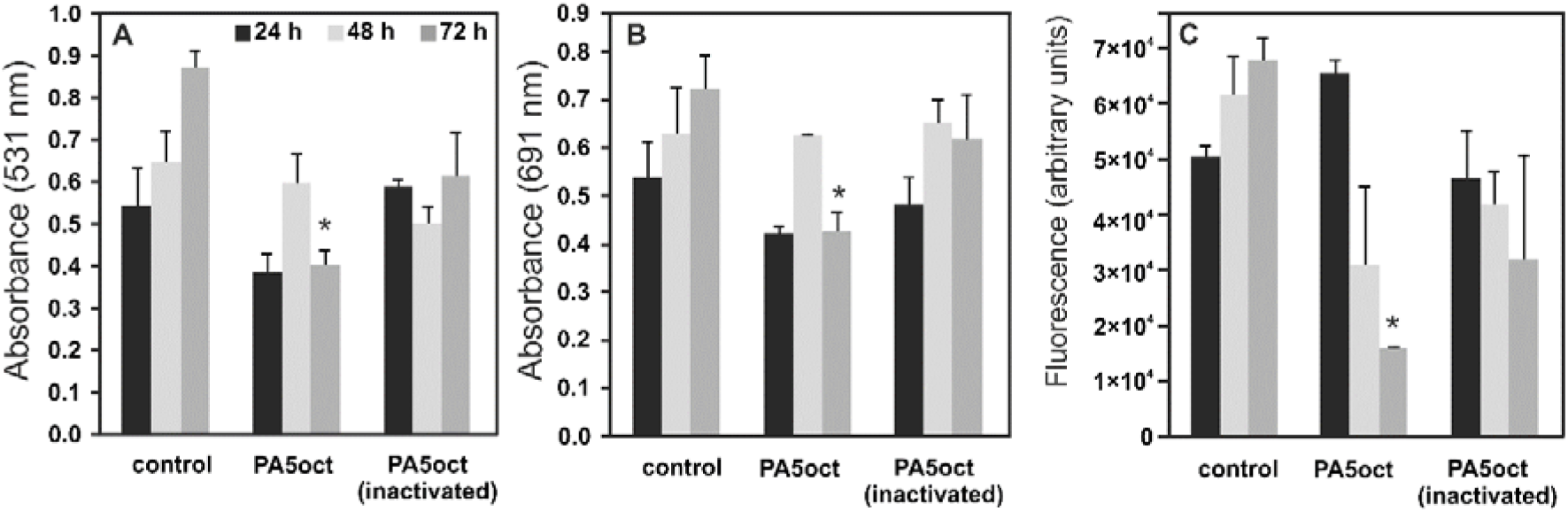
The anti-biofilm effect of PA5oct phage treatment (4 h) on 24, 48 and 72 h PAO1 biofilm formed on Nephrophane membrane. The biomass evaluation by CV staining (A); the level of pyocyanin in growth medium (B); the fluorescence of pyoverdin in growth medium (C). Untreated biofilm was used as control. The results are presented as the means ± SD. Statistical analysis was made by the ANOVA test (denoted p-values < 0.05).

### Phage PA5oct impact on PAO1 biofilm-living population after infection

An important aspect of this study was to evaluate the impact of phage PA5oct’s introduction into a biofilm population. For this purpose, thirty colonies were randomly selected from 24, 48 and 72-hours biofilm after phage treatment and checked for phage susceptibility. Simultaneously, thirty control clones from untreated biofilm were also sampled. A third of phage-exposed clones (10/30) and all control isolates still remained susceptible to PA5oct infection, whereas 20 clones had become resistant. In the next step, isolates were examined for cross resistance to other lytic phages of PAO1 wild type strain (Table S1), recognizing different receptors: Type IV pili-dependent (phiKZ, KTN4, LUZ19) and LPS-dependent (KT28, KTN6, LUZ7, LBL3). Control isolates and PA5oct-sensitive clones taken from the biofilm after phage exposure retain the same phage typing pattern as PAO1 wild type (PA5oct resistant type 0). In total, six different phage typing patterns (0-5) could be distinguished (Table 2).

**Table 2.** Phage typing of PA5oct clones obtained during biofilm treatment

| Bacterial clones | Susceptibility to phage infection |  |  |  |  |  |  |  | Number of isolates |
| --- | --- | --- | --- | --- | --- | --- | --- | --- | --- |
|  | LPS/pili | LPS-dependent |  |  |  | Pili-dependent |  |  |  |
|  | PA5oct | LBL3 | KT28 | KTN6 | LUZ7 | KTN4 | phiKZ | LUZ19 |  |
| Control planktonic PAO1 | + | + | + | + | + | + | + | + | 1 |
| Control biofilm PAO1 | + | + | + | + | + | + | + | + | 30 |
| PA5oct resistant type 0 | + | + | + | + | + | + | + | + | 10 |
| PA5oct resistant type 1 | - | - | + | + | + | - | + | + | 1 |
| PA5oct resistant type 2 | - | - | + | + | + | - | - | + | 2 |
| PA5oct resistant type 3 | - | + | - | - | + | - | + | + | 2 |
| PA5oct resistant type 4 | - | + | - | - | + | + | + | + | 5 |
| PA5oct resistant type 5 | - | - | - | - | + | + | + | + | 10 |

(+) sensitive to phage infection; (-) resistant to phage infection;

In most of the cases, the cross-resistance patterns were related to *Pbunavirus* phages (*Myoviridae*), which primarily target the LPS structure (type 5). However, some PA5oct-resistant clones were resistant to LBL3 phage (*Pbunavirus*) and Type IV pili-dependent giant phages (phiKZ and KTN4). These clones were still susceptible to phages KT28 and KTN6. Three mutants exhibiting resistance to KTN4 phage remained susceptible to its (genomically) closely related phiKZ counterpart (> 99% genome-wide DNA homology) (Danis-Wlodarczyk *et al.*, 2016). These results indicate that PA5oct requires two different receptors for an effective infection. Interestingly, no cross-resistance was observed to the podovirus LUZ7 or other LPS-dependent phages. Similar pattern with respect to the podovirus LUZ19 and the phiKZ-like phages despite their dependence on the same bacterial surface macromolecules for infection (Type IV pili).

### Emerging phage PA5oct-resistant clones showed a reduced virulence

To evaluate the correlation between phage receptor modification in emerging phage-resistant population and the principal virulence factors of these clones, we compared LPS patterns, twitching motility (Type IV pili dependent), growth rate, and *in vivo* pathogenicity in *Galleria mellonella* model (Table 3).

**Table 3.** The virulence features of *P. aeruginosa* clones after PA5oct treatment.

| Bacterial clones | Name | LPS pattern | Growth rate [OD<sub>590</sub> / 24 h] | Larvae survival rate [%] (18/24/48/72 h) | Twitching motility [mm] | Phage DNA presence (PCR) |
| --- | --- | --- | --- | --- | --- | --- |
| control planktonic | PAO1 | S | > 1.5 | 10/0/0/0 | 24.2 ± 1.3 | - |
| control biofilm | K72A | S | > 1.5 | 30/0/0/0 | 23 ± 2.0 | - |
| control biofilm | K72B | S | > 1.5 | 30/0/0/0 | 22.9 ± 1.9 | - |
| control biofilm | K72C | S | > 1.5 | 10/0/0/0 | 22.8 ± 2.3 | - |
| control biofilm | K72D | S | > 1.5 | 5/0/0/0 | 22.7 ± 1.5 | - |
| control biofilm | K72E | S | > 1.5 | 15/0/0/0 | 23.5 ± 1.3 | - |
| control biofilm | K72F | S | > 1.5 | 0/0/0/0 | 23.8 ± 1.2 | - |
| control biofilm | K72G | S | > 1.5 | 5/0/0/0 | 23.7 ± 1.6 | - |
| control biofilm | K72H | S | > 1.5 | 35/0/0/0 | 23.3 ± 2.1 | - |
| control biofilm | K72I | S | > 1.5 | 15/0/0/0 | 23.8 ± 1.1 | - |
| control biofilm | K72J | S | > 1.5 | 25/0/0/0 | 23.2 ± 1.5 | - |
| PA5oct resistant type 0 | 24F | S | < 1.0 | 15/0/0/0 | 18.2 ± 0.9* | - |
| PA5oct resistant type 0 | 48B | S | > 1.5 | 20/10/0/0 | 18.5 ± 1.0* | - |
| PA5oct resistant type 0 | 48C | S | > 1.5 | 80/35/0/0** | 19.3 ± 1.3* | - |
| PA5oct resistant type 0 | 48F | S | > 1.5 | 30/0/0/0 | 15.7 ± 0.8* | - |
| PA5oct resistant type 0 | 48G | S | > 1.5 | 80/40/15/0** | 19 ± 0.8* | - |
| PA5oct resistant type 0 | 72B | S | > 1.5 | 30/25/10/0** | 18.4 ± 1.3* | - |
| PA5oct resistant type 0 | 72C | S | < 1.0 | 80/55/10/0** | 15.5 ± 1.3* | + |
| PA5oct resistant type 0 | 72E | S | > 1.5 | 55/25/0/0** | 14.6 ± 0.7* | - |
| PA5oct resistant type 0 | 72F | S | > 1.5 | 20/0/0/0 | 18.3 ± 0.8* | - |
| PA5oct resistant type 0 | 72G | S | > 1.5 | 30/15/0/0 | 18.5 ± 1.1* | - |
| PA5oct resistant type 1 | 48H | S | < 1.0 | 100/95/75/60** | 5.5 ± 1.3* | + |
| PA5oct resistant type 2 | 24C | S | < 1.0 | 75/60/55/35** | 15.4 ± 1.0* | + |
| PA5oct resistant type 2 | 48A | S | < 1.0 | 70/65/30/0** | 17.5 ± 1.8* | + |
| PA5oct resistant type 3 | 48E | S | < 1.0 | 100/45/0/0** | 19.1 ± 1.3* | + |
| PA5oct resistant type 3 | 72H | S | < 1.0 | 95/55/10/0** | 13.6 ± 1.6* | + |
| PA5oct resistant type 4 | 24A | S | < 1.0 | 50/15/15/15** | 16.7 ± 2.0* | + |
| PA5oct resistant type 4 | 24D | S | > 1.5 | 75/60/40/20** | 11 ± 2.3* | + |
| PA5oct resistant type 4 | 48I | S | < 1.0 | 85/60/20/0** | 13.2 ± 0.6* | + |
| PA5oct resistant type 4 | 48J | S | < 1.0 | 80/60/15/15** | 15.4 ± 1.2* | + |
| PA5oct resistant type 4 | 72I | S | < 1.0 | 55/50/5/0** | 14 ± 1.1* | + |
| PA5oct resistant type 5 | 24B | S | < 1.0 | 85/80/40/40** | 16.5 ± 1.6* | + |
| PA5oct resistant type 5 | 24E | S | < 1.0 | 75/70/55/30** | 15.4 ± 1.6* | + |
| PA5oct resistant type 5 | 24G | S | < 1.0 | 75/70/65/35** | 12.5 ± 2.1* | + |
| PA5oct resistant type 5 | 24H | S | < 1.0 | 95/75/45/35** | 12.2 ± 1.3* | + |
| PA5oct resistant type 5 | 24I | S | > 1.5 | 100/80/55/35** | 13.3 ± 0.8* | + |
| PA5oct resistant type 5 | 24J | S | > 1.5 | 85/70/50/30** | 14.5 ± 1.7* | + |
| PA5oct resistant type 5 | 48D | S | < 1.0 | 85/45/15/0** | 14.4 ± 0.7* | + |
| PA5oct resistant type 5 | 72A | S | < 1.0 | 95/70/0/0** | 12.5 ± 1.7* | + |
| PA5oct resistant type 5 | 72D | S | > 1.5 | 100/75/15/0** | 13.2 ± 0.9* | + |
| PA5oct resistant type 5 | 72J | S | < 1.0 | 95/80/10/0** | 22.8 ± 1.3* | + |

(\*) significantly lower TM zone compared to control clones (One way ANNOVA, P < 0.05)
(\*\*) significantly higher *G. mellonella* larvae survival rate compared to infected by control clones (Log-rank (Mantel-Cox), P < 0.05.

The analysis of LPS extracted from PA5oct phage-resistant types (type 1 - 5) shows no changes compared to both – PA5oct-susceptible mutants (resistance type 0) and biofilm-derived controls (Fig. S1). In the context of these results, cross-resistance to LPS-dependent (KT28, KTN6, LBL3, LUZ7) bacteriophages is an interesting observation. The evaluation of Type IV pili-dependent twitching motility revealed a significant reduction among all the phage-exposed bacteria tested (including resistance type 0), compared to control isolates. Changes in Type IV pili expression may be an explanation for the cross-resistance to pili-dependent phages (KTN4, phiKZ) occurring in 5 out of 20 PA5oct-resistant isolates (Table 3).

The evaluation of growth rate revealed that 8 out of 10 selected PA5oct-susceptible clones (resistance type 0) remained similar to the growth of parental PAO1 and to 30 biofilm-derived controls (OD_590_ > 1.5 after 24h) (Table 3). Among PA5oct-resistant clones 16 out of 20 showed a steep decrease of the growth rate (OD_590_ < 1.0 after 24h) (Table 3, Fig S2).

The impact on the biological features listed above hints towards a modification of the bacterial virulence, which could prove to be a key strategic advantage for the use of PA5oct in therapeutic settings. To validate this observation, we examined the virulence *in vivo*, using the *G. mellonella* infection model (Table 3, Fig. 4, data for remaining clones in supplementary Fig. S2). Using equal doses of bacteria (10 µl of 10^3^ cfu/ml) 25 out of 30 tested clones showed a significant decrease in virulence (P < 0.05), including 5 PA5oct-susceptible clones (resistance type 0). We were finally able to distinguish four distinct phenotypic patterns in PAO1 clones: fast-growing with a high virulence (Fig. 4A/B); slow-growing with a high virulence (Fig. 4C/D); fast-growing with a low virulence (Fig. 4E/F), and slow-growing with a low virulence (Fig. 4G/H). The relationship between growth rate and virulence of all phage-treated bacterial mutants are summarized in Table 3 and Fig. S2.

**Figure 4.**
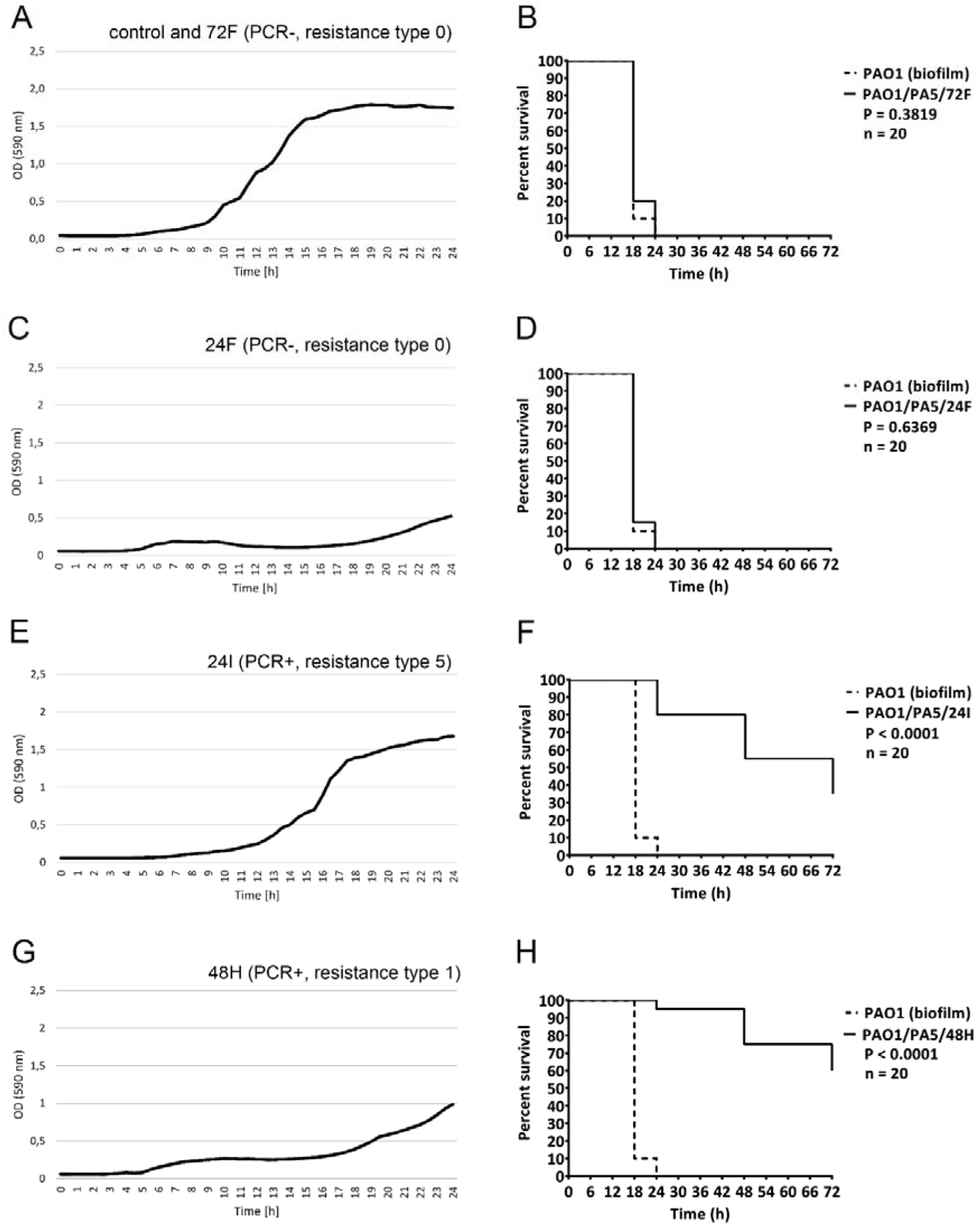
Phenotypic patterns in PAO1 clones treated with PA5oct. Growth curves (A, C, E, G) of selected bacterial clones combined with a survival of infected *G. mellonella* (B, D, F, H) for PA5oct-resistant clones. Fast-growing with a high virulence (control and 72F; Fig. 4A/B); slow-growing with a high virulence (24F; Fig. 4C/D); fast-growing with a low virulence (24I; Fig. 4E/F), and slow-growing with a low virulence (48H; Fig. 4G/H). Data for all clones are presented in supplementary Fig. S2.

### The reduced virulence in PA5oct-resistant clones is correlated with the persistence of phage in bacterial population

The analysis of 30 phage-treated clones indicates that the reduced virulence has emerged as a consequence of PA5oct infection of the PAO1 population. Moreover, slow-growing isolates may suggest a phage propagation event in the bacterial culture. Therefore, a PCR assay targeting the unique gene encoding the major head subunit precursor in PA5oct was implemented to confirm the presence of phage DNA within bacterial population. All PA5oct-resistant isolates and one still sensitive (72C) were PCR-positives (Table 3) and all those clones were less virulent compared to wild type PAO1. Moreover, half of the clones that were still sensitive to phage infection became less virulent as well. A high prevalence of phage maintenance within resistant bacterial cells might suggest prolonged phage propagation or pseudolysogeny/carrier state event.

### The presence of PA5oct in PAO1 population induce pro-inflammatory response in monocytes

Given the results listed above, we were interested to investigate the possible influence of PA5oct infection on pro-inflammatory features of *Pseudomonas* population, which would further support the antimicrobial activity obtained in the ASL and larvae models. Toll-like Receptors (TLRs) present on phagocytic cells serve as Pattern Recognition Receptors (PRRs) and play a crucial role in the proper functioning of the innate immune system. Therefore, we selected 10 PA5oct-resistant clones and determined the TLR stimulation profile of THP1-XBlue™ monocytes line when treated with bacteria culture filtrates. The results correlated perfectly to our *in vivo* data, in which the same mutants exhibiting a low virulence and PCR-positive, gained pro-inflammatory features, strongly stimulated the monocyte culture, relative to PAO1 isolate controls (Fig. S3).

## Discussion

The principal aim of this study was to assess the impact of the jumbo phage PA5oct’s infection on *P. aeruginosa* PAO1 population. An integral element in the establishment of phage PA5oct biology is phage receptor analysis. Knock-out strains and cross resistance experiments revealed a dependence on LPS and Type IV pili receptor. This versatility can generally explain the broad spectrum of lytic activity of jumbo phages. In our case, the distinct host range between both strain panels is noteworthy. In the reference panel of *P. aeruginosa* (BCCM / LMG), containing 43 strains (25 isolated from patients with cystic fibrosis) (De Soyza *et al.*, 2013), only 24% of isolates showed the susceptibility to PA5oct phage (6 from CF patients) (Cullen *et al.*, 2015). Concurrently, phage PA5octis able to infect 40% of isolates from our standard clinical strain panel. This is probably due to the specific phenotype of long-term CF-related strains, including reduced virulence patterns compared to environmental strains (modified LPS structure, lower expression of Type IV pili and flagellum and a decrease of production intensity of alginate, pyocyanin, pyoverdine and elastase) (Cigana *et al.*, 2009; Bradbury *et al.*, 2011; Lorè *et al.*, 2012).

We used the ASL model to mimic natural conditions of *Pseudomonas* epithelium infection *in vitro*. The experiments confirmed that PA5oct can efficiently access and infect planktonic, mucus embedded and cell-adhered bacteria, but remained dependent on the strain used, as previously observed for giant *Pseudomonas* KTN4 phage (Worlitzsch *et al.*, 2002; Danis-Wlodarczyk *et al.*, 2016). According to a study by Worlitzsch *et al.* (Worlitzsch *et al.*, 2002) using CuFi-1 epithelial cell line, *P. aeruginosa* does not interact the CF epithelium directly, but rather gets embedded in mucus plugs formed in the airways.

*P. aeruginosa* strains entering the human respiratory tract usually express the entire arsenal of virulence factors including flagellum, LPS, Type IV fimbriae or alginate (Prince, 1992). It enables bacteria to successfully adhere to and invade respiratory epithelium. Simultaneously, the exposition of bacterial surface structures exposes them to potential phage infection (phages usually target LPS and Type IV fimbriae). Under natural conditions, bacterial infection stimulate the pro-inflammatory response of immune and epithelial cells, which results in an increased production of antibacterial peptides, cytokines and recruitment of phagocytic cells (Saiman *et al.*, 2001; Dechecchi *et al.*, 2007, 2008; Dean *et al.*, 2011; Dalcin and Ulanova, 2013). In the experimental case of epithelial NuLI-1 and CuFI-1 cell lines (Zabner *et al.*, 2003), a significant decrease in bacterial cell counts demonstrated in ASL studies may be linked to the synergistic action of PA5oct phage and antibacterial peptides produced by the epithelial cell lines tested. Moreover, the release of pro-inflammatory compounds from *Pseudomonas* population infected with this jumbo phage resulted in increased stimulation in monocytic cells enhancing the clearance mechanisms of innate immune system. This further supports our results in both ASL and *in vivo* model.

From the perspective of sessile cells, the infection by phages is limited due to the presence of biofilm matrix, the modification of bacterial surface receptors and reduced metabolic activity (Labrie *et al.*, 2010; Abedon, 2017). Apart from being able to diffuse in a dense airway mucus, PA5oct could also get an access to bacterial cells hidden in a mature 72-hour biofilm. This was verified by our two complementary assays (permeability assay and Nephrophane membrane biofilm assay). That observation is in contradiction with the general view that phages have the greatest activity against immature biofilms composed mostly of metabolically active cells (Azeredo and Sutherland, 2008; Abedon, 2016). In the case of PA5oct, biofilm biomass reduction occurred only as a result of phage-mediated cell lysis, whereas biofilm matrix was degraded by both active and inactivated phages (noticeable increase of TSA diffusion trough the Nephrophane membrane). Such result might suggest that PA5oct phage destroys the biofilm not only by unsealing its structure after tightly packed cells lysis, but it might also use exopolysaccharide depolymerases (potential candidates gp162 and gp205), which are probably responsible for matrix disruption (Pires *et al.*, 2016; Olszak *et al.*, 2017). The introduction of PA5oct phage into the biofilm population of *P. aeruginosa* triggered the emergence of clones resistant to this virus. The PA5oct-resistant isolates subjected to typing with a panel of phages recognising different receptors, showed that the phenotypic changes under the influence of PA5oct pressure went beyond the modulation of a single receptor. PA5oct-resistant isolates showed a cross-resistance to phages belonging to different taxonomic units, and the typing patterns were diverse, suggesting genomic rearrangements in the bacterial genome which impact multiple receptors (Le *et al.*, 2014; Shen *et al.*, 2018). These changes in the genome of the host are currently under investigation by our team.

The formation of phage-resistant bacterial variants is often thought to increase virulence, calling for skepticism on the implementation of phage therapy. Although phage-related phenotypic changes in bacteria decrease the efficacy of phage therapy as a standalone therapy, it turns out that in practice the occurring phenotypic modifications usually lead to a diminished bacterial virulence. Consequently, the *Pseudomonas* population that survived the phage treatment/infection became sensitive to the immune system that effectively removes the pathogenic agent. Our findings were confirmed in the *G. mellonella* infection model, but also reported by others in murine model (Roach *et al.*, 2017).

An often neglected phenomenon is the occurrence of pseudolysogeny/carrier stage, generally regarded as a temporary stage of phage particle dormancy. However, it appears that the presence of a phage episome inside the host cell influences its phenotype and contributes the cross-resistance to other bacteriophages as well (Cenens et al., 2015; Latino *et al.*, 2016; Argov *et al.*, 2017). The jumbo phages are known to easily undergo episome formation as was previously reported for phiKZ phage (Krylov *et al.*, 2012). In this study, we indicated the presence of PA5oct DNA in isolated resistant-clones and linked it to modified virulence of the PAO1 population carrying the phage DNA. A significant survival improvement was seen in larvae infected with those isolates. These strains became significantly less pathogenic *in vivo* compared to wild-type PAO1 and to PCR-negative phage-resistant mutants. This effect could potentially be directly or indirectly induced by the expression of specific PA5oct genes. Taking into account that the prevalence of PCR-positive clones was relatively high (21/30 strains) we may conclude that this specific jumbo phage PA5oct efficiently eradicates sensitive *P. aeruginosa* cells both planktonic and sessile, while at the same time selects for a primarily non-virulent pseudolysogenic/carrier resistant population.

Owing to its unusual behavior when compared to other known virulent phages, PA5oct proved to be an interesting subject for research in the ecological and evolutionary context. Its anti-biofilm activity and the influence on bacterial virulence factors expression exert a strong selective pressure within the bacterial population. This phenomenon, combined with the long-term persistence of PA5oct in the bacterial population (pseudolysogeny/carrier stage) and the generation of cross-resistance to unrelated species of viruses, is an important item for further research that can contribute to our understanding of co-evolution of bacteria and their phages. The pro-inflammatory activity of bacterial clones infected by phages causing an increased susceptibility to the innate immune system is another interesting aspect which is currently being investigated.

## Experimental procedures

### Bacteriophages propagation and purification

Phage PA5oct was propagated as previously described by Danis-Wlodarczyk (Danis-Wlodarczyk *et al.*, 2015). Phage lysate was purified by 0.45 and 0.22 µm filtration and the incubation with 10% polyethylene glycol 8000 (PEG 8000) – 1 M NaCl according to standard procedures (Ceyssens *et al.*, 2006). Finally, the CsCl-gradient ultracentrifugation was applied (Ceyssens *et al.*, 2008) and phage preparation was dialyzed three times for 30 min against 250 volumes of phage buffer using Slide-A-Lyzer Dialysis Cassettes G2 (Thermo Fisher Scientific Inc, MA, USA). The phage titre was assessed using the double-agar layer technique (Adams, 1959) and purified samples were stored at 4°C in the dark.

### Phage host range and phage receptor analysis

The bacterial susceptibility to phage-mediated lysis was determined by a standard spot test assay, applying serial dilution of phage titer (10^8^-10^3^ pfu/ml) (Kutter, 2009). The phage receptor on bacterial surface was evaluated on PAO1 knock-out mutants deficient in biosynthesis of A-band and B-band O-antigen, flagella, Type IV pili, or alginate production (Table 1). The phage host range was evaluated on 47 CF strains from the Leuven University hospital, Leuven, Belgium collection (Lood *et al.*, in preparation) and compared to other *Pseudomonas* phages listed in Table S1-S2.

### Airway Surface Liquid infection model

For the Airway Surface Liquid (ASL) experiments, two cell lines (NuLi-1; Normal Lung, University of Iowa), isolated from human airway epithelium of normal genotype and CuFi-1; Cystic Fibrosis, University of Iowa, isolated from bronchial epithelium of a CFTR F508del/F508del patient) were used (kindly provided by prof. Zabner, University of Iowa, Iowa City, IA). The ASL model was prepared according to methods described by Zabner (Zabner *et al.*, 2003). The experiment was conducted as previously described (Danis-Wlodarczyk *et al.*, 2016). In short, cell lines were infected with the *P. aeruginosa* PAO1 reference strain (6.2 × 10^7^ cfu/ml), nonCF0038 isolate from burn wound (6.5 × 10^7^ cfu/ml) and CF708 small colony variant (1.0 × 10^6^ cfu/ml) and incubated for 1.5 h at 37°C, 5% CO_2_. Next, a 25 μl volume of the PA5oct phage lysate (6 × 10^8^ pfu/ml) was added to each millicell hanging cell culture insert, which were further incubated for 1.5 h at 37°C, 5% CO_2_. To evaluate the phage efficacy in bacteria eradication, cells after treatment were washed with PBS and CFU counts were calculated from apical washes on LB agar (Sigma-Aldrich). The data was analyzed using the Statistica software package (StatSoft, Tulsa, OK, USA). All values were expressed as mean ± SD and significant differences between variations (denoted p-values < 0.05) were assessed using the Snedecor-Fisher test using one-way ANOVA.

### Biofilm eradication analysis on Nephrophane membrane

The ability of PA5oct phage to degrade biofilm matrix was evaluated by laser interferometry method (Danis-Wlodarczyk *et al.*, 2016). PAO1 biofilm was formed for 72 h at 37°C in Trypticase-Soy broth (TSB, bioMerieux, France), as the level of membrane coverage by biofilm was determined as around 93 % (Fig 2A) after removing of supernant. Next, the biofilm was treated for four hours at 37°C with an intact or UV-inactivated phage suspensions (5 × 10^8^ pfu/ml). After treatment the membrane was washed with saline and examined by interferometry technique. The degradation of biofilm was assessed as the permeability increase of its matrix for low molecular compounds (TSB). The quantitative measurements of TSB diffusion through biofilm structure treated with phage was tested by laser interferometry method (Danis-Wlodarczyk *et al.*, 2015, 2016). The data was analyzed using the Statistica software package (StatSoft, Tulsa, OK, USA). All values obtained at the end of 40 min measurements were expressed as mean ± SD and significant differences between variations (denoted p-values < 0.05) were assessed using the Snedecor-Fisher test using a one-way ANOVA.

Biofilm eradication was also examined by crystal violet (0.004%) staining for 15 min as previously described (Danis-Wlodarczyk *et al.*, 2016) and tested for pyocyanin and pyoverdin/pyochelin levels in the supernatants as described below.

### Isolation of phage resistant clones from treated biofilm

*P. aeruginosa* PAO1 mutants resistant to PA5oct phage were isolated by the controlled infection. In the first step, bacterial suspension in TSB was incubated (37°C) in 96-well peg-lid plate (Nunc, Denmark) for 24, 48 or 72 hours enabling bacteria to form biofilm. Afterwards, the mature biofilm molded on the surface of pegs was immersed in phage PA5oct suspension (10^6^ pfu/ml) for 24 hours. Finally, pegs were washed with PBS buffer to clear away planktonic bacteria and the biofilm population was collected using an ultrasonic bath, and then plated on TSA (bioMerieux) for 24 h at 37°C, to isolate discrete colonies. Ten randomly selected isolates from each time point, were passaged five times on TSA to confirm the stability of genetic changes. Thirty control non-treated biofilm strains were isolated in an analogous manner. The emerging PA5oct phage-resistant clones were verified by spot technique, applying 10^5^ pfu/ml PA5oct phage titer (Kutter, 2009). Moreover, the clones were also tested the same way in terms of the susceptibility to other *Pseudomonas* phages: Type IV pili-dependent (phiKZ, KTN4, LUZ19), LPS-dependent (KT28, KTN6, LUZ7) and a phage with an unverified receptor (LBL3) (Table S1). All tests were performed in triplicate.

### Pyocyanin and pyoverdine production

The level of pyoverdine and pyocyanine production was analyzed on a 72h-biofilm formed on Nephrophane or selected PAO1 mutants cultured (48 h, 37°C) in TSB medium (bioMerieux, France) in a 24-well polystyrene plate (Sarstedt, Germany) (Danis-Wlodarczyk *et al.*, 2015). To investigate the level of pyocyanin production the supernatant absorbance was measured at λ = 695 nm. For pyoverdine production the fluorescence was measured at λex = 392 nm and λem = 460 nm.

### Twitching motility assay

Solid surface bacterial movement dependent on Type IV pili, called “twitching motility” was measured according to Turnbull & Whitchurch (Turnbull and Whitchurch, 2014). Each bacterial suspension was transferred to agar plate using a toothpick by perpendicular stabbing through the agar layer (up to the bottom of the plate) and incubated for 48 h at 37□. After incubation the agar layer was removed and the 0.01% crystal violet was used for visualization of the growth zone. The growth zone results for twitching motility are presented as the mean of ten replicates.

### Lipopolysaccharide structure patterns analysis

Examination of LPS structure pattern was done using slightly modified Marolda method of extraction (Marolda *et al.*, 1990) and SDS-PAGE. Overnight bacterial cultures in TSB medium (bioMerieux, France) were centrifuged and adjusted to OD_600_ = 2.0 in PBS buffer. Bacterial cells were destroyed by boiling in lysis buffer (2% SDS, 4% β-mercaptoethanol, Tris, pH 6.8) and digesting with proteinase K (60°C, 1h). Protein debris were eliminated by incubation (70°C, 15 min.) with equal volume of 90% phenol. LPS containing aqueous phase was recovered using ethyl ether to remove any residual phenol. To assess the concentration of LPS used for electrophoresis, the Purpald method of KDO (3-Deoxy-D-manno-oct-2-ulosonic acid) measurement was used (Lee and Tsai, 1999). Subsequently, the LPS samples were separated by Tricine SDS-PAGE method (14% polyacrylamide gel with 4M urea, 125 V). Finally, after separation, LPS samples were stained with silver according to Tsai & Frasch (Tsai and Frasch, 1982).

### Growth rate measurement

The growth rate of selected isolates was estimated using the indirect method, by measuring the optical density of a 24-hour culture at a 30 min. interval. The 18-hour TSA plate cultures were suspended in PBS with an optical density of 0.5 McF and used to establish a liquid cultures in a 24-well titration plate (starting cfu/ml = 10^6^). A plate with applied cultures was placed inside of a microplate reader (Varioscan Lux, Thermo Scientific) and incubated for 24 hours. The absorbance measurement (λ = 590 nm) was performed automatically every 30 min. after shaking for 30 s. The results of the growth rate measurements were classified in two categories: HIGH (OD_590_ after 24 h > 1.5) and LOW (OD_590_ after 24 h < 1.0).

### TLR stimulation profile in THP1-XBlue™ cell line, NF-kB/AP-1-Reporter Monocytes

The Toll-like receptors (TLRs) stimulation of monocytes, major forms of innate immune sensors, was assessed according to manufacturer protocol using THP1-XBlue™ cell line (InvivoGen, Toulouse France). THP1-XBlue™ cells derive from the human monocytic THP-1 cell line and express an NF-κB- and AP-1-inducible secreted embryonic alkaline phosphatase (SEAP) reporter gene. Upon TLR2, TLR1/2, TLR2/6, TLR4, TLR5 and TLR8 stimulation, THP1-XBlue™ cells activate transcription factors and subsequently the secretion of SEAP which is easily detectable when using QUANTI-Blue™, a medium that turns purple/blue in the presence of SEAP. The results were analyzed using a microplate reader (Varioscan Lux, Thermo Scientific).

### Galleria mellonella *larvae infection model*

The virulence of PAO1 mutants resistant to PA5oct phage was tested *in vivo* in *G. mellonella* infection model, previously described by Cullen et al. (Cullen *et al.*, 2015). The wax moth larvae were sorted by size and weight and acclimated for one week at 15°C. The larvae were infected by the injection into the hindmost proleg, of 10 µl of bacterial suspension (10^3^ cfu/ml) giving 10 cells per larvae. Further, the larvae were incubated at 37°C for 72h, and their viability was checked after 8, 18, 24, 48 and 72 hours post injection. The two types of control were made: in negative control larvae were injected with 10µl of sterile PBS buffer, while the positive control consisted of wild PAO1 strain infection. The graphical presentation of the survival curves was prepared using GraphPad Prism 6.0 (GraphPad Software Inc., La Jolla, USA). The log-rank Mantel-Cox test was used for statistical analysis (*P*-values < 0.05 were regarded as significant).

### PA5oct phage DNA detection in bacterial clones

The presence of PA5oct genome within the cells of PAO1 resistant mutants was evaluated using a standard polymerase chain reaction (PCR). The DNA was isolated using PureLink Genomic DNA mini kit (Invitrogen, Thermo Scientific). On the basis of the full sequence of PA5oct genome, primers flanking a fragment of the structural gene (major head subunit precursor) were designed (F: 5’-GATACATACCCTACGTGTTCGTTATG-3’ and R: 5’- GCACCGTTACCCAGCGAGTTAG). The PCR was carried out under the optimized conditions: initialization (95°C / 5 min), 30 cycles of denaturation (95°C / 30 s), annealing (56.4°C / 1 min) and elongation (72°C / 1 min 10 s) followed by final elongation (72°C / 10 min). The resulting 872 bp reaction product was visualized by standard agarose gel electrophoresis (1% agarose, 1X TBE buffer, 95 V/cm / 45 min.). The positive control was a purified PA5oct phage preparation and the negative control was a PAO1 strain (Fig. S4).

## Acknowledgements

This study was supported by research grant 2012/04/M/NZ6/00335 and 2015/18/M/NZ6/00413 of National Science Centre, Poland. RL is supported by a GOA grant entitled “Phage Biosystems” from the KU Leuven. CL is supported by an SB PhD fellowship from FWO Vlaanderen (1S64718N).

## Supplementary materials

Table S1. Main features of *Pseudomonas* phages used in the study.

Table S2. Phage activity comparison of six virulent phages on *P. aeruginosa* strains from the University hospital of Leuven, Leuven, Belgium collection.

Figure S1. LPS profiles of PA5-phage-resistant clones analyzed in 14% polyacrylamide/tricine-SDS gels.

Figure S2. Comparison of the growth rate, virulence (on *G. mellonella* model) and phage resistance pattern of 30 clones of *P. aeruginosa* PAO1 isolated as a result of controlled PA5oct phage infection.

Figure S3. THP-1 X-blue monocyte response to PA5-phage-resistant clones post-culture medium stimulation.

Figure S4. PCR analysis targeting the major head subunit precursor gene in selected PA5oct resistant clones. M-mass marker.

## References

Abedon, S.T. (2016) Bacteriophage exploitation of bacterial biofilms: phage preference for less mature targets? FEMS Microbiol. Lett. 363: fnv246.

Abedon, S.T. (2017) Phage “delay” towards enhancing bacterial escape from biofilms: a more comprehensive way of viewing resistance to bacteriophages. AIMS Microbiol. 3: 186–226.

Adams, M.H. (1959) Enumeration of phage particles. In, Adams, M.H. (ed), Bacteriophages. Interscience Publishers, New York, pp. 27–30.

Argov, T., Azulay, G., Pasechnek, A., Stadnyuk, O., Ran-Sapir, S., Borovok, I., et al. (2017) Temperate bacteriophages as regulators of host behavior. Curr. Opin. Microbiol. 38: 81–87.

Azeredo, J. and Sutherland, I. (2008) The use of phages for the removal of infectious biofilms. Curr. Pharm. Biotechnol. 9: 261–266.

Bradbury, R.S., Reid, D.W.E.C., Inglis, T.J.J., and Champion, A.C. (2011) Decreased virulence of cystic fibrosis *Pseudomonas aeruginosa* in *Dictyostelium discoideum*. Microbiol. Immunol. 55: 224–230.

Ceyssens, P.-J., Lavigne, R., Mattheus, W., Chibeu, A., Hertveldt, K., Mast, J., et al. (2006) Genomic analysis of *Pseudomonas aeruginosa* phages LKD16 and LKA1: establishment of the KMV subgroup within the T7 supergroup. J. Bacteriol. 188: 6924–6931.

Ceyssens, P.-J., Mesyanzhinov, V., Sykilinda, N., Briers, Y., Roucourt, B., Lavigne, R., et al. (2008) The genome and structural proteome of YuA, a new *Pseudomonas aeruginosa* phage resembling M6. J. Bacteriol. 190: 1429–1435.

Ceyssens, P.-J., Minakhin, L., Van den Bossche, A., Yakunina, M., Klimuk, E., Blasdel, B., et al. (2014) Development of giant bacteriophage phiKZ is independent of the host transcription apparatus. J. Virol. 88: 10501–15010.

Chaikeeratisak, V., Nguyen, K., Khanna, K., Brilot, A.F., Erb, M.L., Coker, J.K.C., et al. (2017) Assembly of a nucleus-like structure during viral replication in bacteria. Science (80-.). 355: 194–197.

Cigana, C., Curcurù, L., Leone, M.R., Ieranò, T., Lorè, N.I., Bianconi, I., et al. (2009) *Pseudomonas aeruginosa* exploits lipid A and muropeptides modification as a strategy to lower innate immunity during cystic fibrosis lung infection. PLoS One 4: e8439.

Cullen, L., Weiser, R., Olszak, T., Maldonado, R., Moreira, A., Slachmuylders, L., et al. (2015) Phenotypic characterization of an international *Pseudomonas aeruginosa* reference panel: strains of cystic fibrosis (CF) origin show less in vivo virulence than non-CF strains. Microbiology 161: 1961–1977.

Dalcin, D. and Ulanova, M. (2013) The role of human beta-defensin-2 in Pseudomonas aeruginosa pulmonary infection in cystic pibrosis patients. Infect. Dis. Ther. 2: 159–166.

Danis-Wlodarczyk, K., Olszak, T., Arabski, M., Wasik, S., Majkowska-Skrobek, G., Augustyniak, D., et al. (2015) Characterization of the newly isolated lytic bacteriophages KTN6 and KT28 and their efficacy against *Pseudomonas aeruginosa* biofilm. PLoS One 10: e0127603.

Danis-Wlodarczyk, K., Vandenheuvel, D., Jang, H. Bin, Briers, Y., Olszak, T., Arabski, M., et al. (2016) A proposed integrated approach for the preclinical evaluation of phage therapy in *Pseudomonas* infections. Sci. Rep. 6: 28115.

Dean, S.N., Bishop, B.M., and van Hoek, M.L. (2011) Susceptibility of *Pseudomonas aeruginosa* biofilm to alpha-helical peptides: D-enantiomer of LL-37. Front. Microbiol. 2: 128.

Dechecchi, M.C., Nicolis, E., Bezzerri, V., Vella, A., Colombatti, M., Assael, B.M., et al. (2007) MPB-07 reduces the inflammatory response to *Pseudomonas aeruginosa* in cystic fibrosis bronchial cells. Am. J. Respir. Cell Mol. Biol. 36: 615–624.

Dechecchi, M.C., Nicolis, E., Norez, C., Bezzerri, V., Borgatti, M., Mancini, I., et al. (2008) Anti-inflammatory effect of miglustat in bronchial epithelial cells. J. Cyst. Fibros. 7: 555–565.

Drulis-Kawa, Z., Olszak, T., Danis, K., Majkowska-Skrobek, G., and Ackermann, H.-W. (2014) A giant *Pseudomonas* phage from Poland. Arch. Virol. 159: 567–572.

Fokine, A., Battisti, A.J., Bowman, V.D., Efimov, A. V., Kurochkina, L.P., Chipman, P.R., et al. (2007) Cryo-EM Study of the *Pseudomonas* bacteriophage φKZ. Structure 15: 1099–1104.

Forti, F., Roach, D.R., Cafora, M., Pasini, M.E., Horner, D.S., Fiscarelli, E. V., et al. (2018) Design of a broad-range bacteriophage cocktail that reduces *Pseudomonas aeruginosa* biofilms and treats acute infections in two animal models. Antimicrob. Agents Chemother. 62: e02573–17.

Galtier, M., De Sordi, L., Sivignon, A., de Vallée, A., Maura, D., Neut, C., et al. (2017) Bacteriophages targeting adherent invasive *Escherichia coli* strains as a promising new treatment for Crohn’s disease. J. Crohn’s Colitis 11: 840–847.

Hendrix, R.W. (2009) Jumbo bacteriophages. Curr. Top. Microbiol. Immunol. 328: 229–240.

Krylov, V., Shaburova, O., Krylov, S., and Pleteneva, E. (2012) A genetic approach to the development of new therapeutic phages to fight *Pseudomonas aeruginosa* in wound infections. Viruses 5: 15–53.

Krylov, V.N., Dela Cruz, D.M., Hertveldt, K., and Ackermann, H.-W. (2007) “φKZ-like viruses”, a proposed new genus of myovirus bacteriophages. Arch. Virol. 152: 1955–1959.

Kutter, E. (2009) Phage host range and efficiency of plating. Methods Mol. Biol. 501: 141–149.

Labrie, S.J., Samson, J.E., and Moineau, S. (2010) Bacteriophage resistance mechanisms. Nat. Rev. Microbiol. 8: 317–327.

Latino, L., Midoux, C., Hauck, Y., Vergnaud, G., and Pourcel, C. (2016) Pseudolysogeny and sequential mutations build multiresistance to virulent bacteriophages in *Pseudomonas aeruginosa*. Microbiol. (United Kingdom) 162: 748–763.

Le, S., Yao, X., Lu, S., Tan, Y., Rao, X., Li, M., et al. (2014) Chromosomal DNA deletion confers phage resistance to *Pseudomonas aeruginosa*. Sci. Rep. 4: 4738.

Lecoutere, E., Ceyssens, P.J., Miroshnikov, K.A., Mesyanzhinov, V. V., Krylov, V.N., Noben, J.P., et al. (2009) Identification and comparative analysis of the structural proteomes of φKZ and EL, two giant *Pseudomonas aeruginosa* bacteriophages. Proteomics 9: 3215–3219.

Lee, C.H. and Tsai, C.M. (1999) Quantification of bacterial lipopolysaccharides by the purpald assay: Measuring formaldehyde generated from 2-keto-3-deoxyoctonate and heptose at the inner core by periodate oxidation. Anal. Biochem. 267: 161–168.

Lorè, N.I., Cigana, C., De Fino, I., Riva, C., Juhas, M., Schwager, S., et al. (2012) Cystic fibrosis-niche adaptation of *Pseudomonas aeruginosa* reduces virulence in multiple infection hosts. PLoS One 7: e35648.

Marolda, C.L., Welsh, J., Dafoe, L., and Valvano, M.A. (1990) Genetic analysis of the O7-polysaccharide biosynthesis region from the *Escherichia coli* O7:K1 strain VW187. J. Bacteriol. 172: 3590–3599.

Mesyanzhinov, V. V, Robben, J., Grymonprez, B., Kostyuchenko, V.A., Bourkaltseva, M. V, Sykilinda, N.N., et al. (2002) The genome of bacteriophage φKZ of *Pseudomonas aeruginosa*. J. Mol. Biol. 317: 1–19.

Monson, R., Foulds, I., Foweraker, J., Welch, M., and Salmond, G.P.C. (2011) The *Pseudomonas aeruginosa* generalized transducing phage phiPA3 is a new member of the phiKZ-like group of “jumbo” phages, and infects model laboratory strains and clinical isolates from cystic fibrosis patients. Microbiology 157: 859–67.

Olszak, T., Shneider, M.M., Latka, A., Maciejewska, B., Browning, C., Sycheva, L. V., et al. (2017) The O-specific polysaccharide lyase from the phage LKA1 tailspike reduces *Pseudomonas* virulence. Sci. Rep. 7: 16302.

Olszak, T., Zarnowiec, P., Kaca, W., Danis-Wlodarczyk, K., Augustyniak, D., Drevinek, P., et al. (2015) *In vitro* and *in vivo* antibacterial activity of environmental bacteriophages against *Pseudomonas aeruginosa* strains from cystic fibrosis patients. Appl. Microbiol. Biotechnol. 99: 6021–6033.

Pires, D.P., Oliveira, H., Melo, L.D.R., Sillankorva, S., and Azeredo, J. (2016) Bacteriophage-encoded depolymerases: their diversity and biotechnological applications. Appl. Microbiol. Biotechnol. 100: 2141–2151.

Pirnay, J.-P., Verbeken, G., Ceyssens, P.-J., Huys, I., De Vos, D., Ameloot, C., and Fauconnier, A. (2018) The Magistral phage. Viruses 10: 64.

Pirnay, J.-P., De Vos, D., Cochez, C., Bilocq, F., Vanderkelen, A., Zizi, M., et al. (2002) *Pseudomonas aeruginosa* displays an epidemic population structure. Environ. Microbiol. 4: 898–911.

Prince, A. (1992) Adhesins and receptors of *Pseudomonas aeruginosa* associated with infection of the respiratory tract. Microb. Pathog. 13: 251–260.

Roach, D.R., Leung, C.Y., Henry, M., Morello, E., Singh, D., Di Santo, J.P., et al. (2017) Synergy between the host immune system and bacteriophage is essential for successful phage therapy against an acute respiratory pathogen. Cell Host Microbe 22: 38–47.e4.

Saiman, L., Tabibi, S., Starner, T.D., San Gabriel, P., Winokur, P.L., Hong Peng Jia, et al. (2001) Cathelicidin peptides inhibit multiply antibiotic-resistant pathogens from patients with cystic fibrosis. Antimicrob. Agents Chemother. 45: 2838–2844.

Schneider, C.A., Rasband, W.S., and Eliceiri, K.W. (2012) NIH Image to ImageJ: 25 years of image analysis. Nat. Methods 9: 671–675.

Shen, M., Zhang, H., Shen, W., Zou, Z., Lu, S., Li, G., et al. (2018) *Pseudomonas aeruginosa* MutL promotes large chromosomal deletions through non-homologous end joining to prevent bacteriophage predation. Nucleic Acids Res.

De Soyza, A., Hall, A.J., Mahenthiralingam, E., Drevinek, P., Kaca, W., Drulis-Kawa, Z., et al. (2013) Developing an international *Pseudomonas aeruginosa* reference panel. Microbiologyopen 2: 1010–1023.

Tsai, C.M. and Frasch, C.E. (1982) A sensitive silver stain for detecting lipopolysaccharides in polyacrylamide gels. Anal. Biochem. 119: 115–119.

Turnbull, L. and Whitchurch, C.B. (2014) Motility assay: twitching motility. In, Filloux, A. and Ramos, J.-L. (eds), Pseudomonas Methods and Protocols. Humana Press Inc., New York, pp. 73–86.

Wittebole, X., De Roock, S., and Opal, S.M. (2014) A historical overview of bacteriophage therapy as an alternative to antibiotics for the treatment of bacterial pathogens. Virulence 5: 226–235.

Worlitzsch, D., Tarran, R., Ulrich, M., Schwab, U., Cekici, A., Meyer, K.C., et al. (2002) Effects of reduced mucus oxygen concentration in airway *Pseudomonas infections* of cystic fibrosis patients. J. Clin. Invest. 109: 317–325.

Yuan, Y. and Gao, M. (2017) Jumbo bacteriophages: An overview. Front. Microbiol. 8:.

Zabner, J., Karp, P., Seiler, M., Phillips, S.L., Mitchell, C.J., Saavedra, M., et al. (2003) Development of cystic fibrosis and noncystic fibrosis airway cell lines. Am. J. Physiol. Cell. Mol. Physiol. 284: L844–L854.

