## Supplementary figures and images for "*Pseudomonas aeruginosa* PA5oct jumbo phage impacts planktonic and biofilm population and reduces its host virulence"

### Supplemental Fig. S1

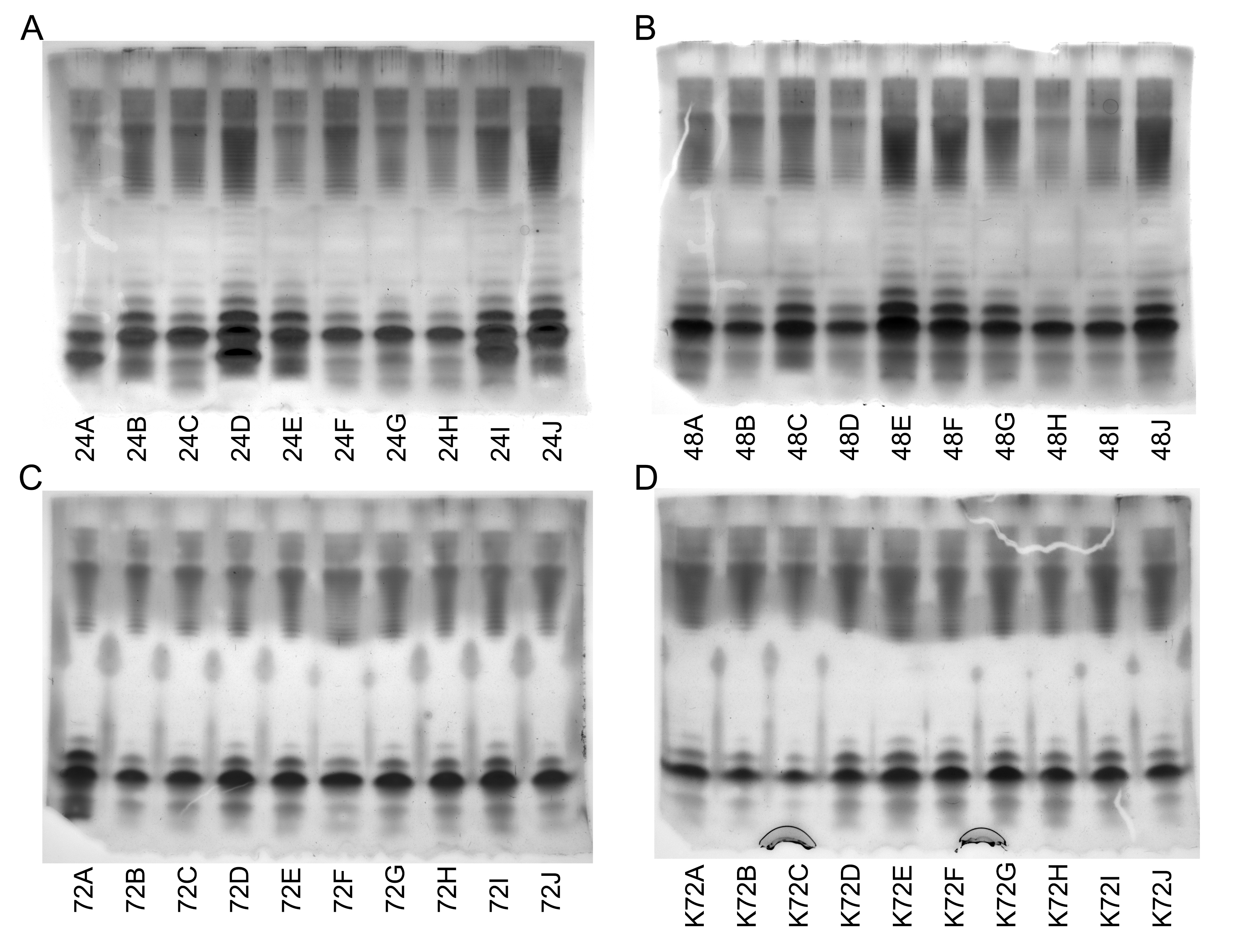

### Supplemental Fig. S2

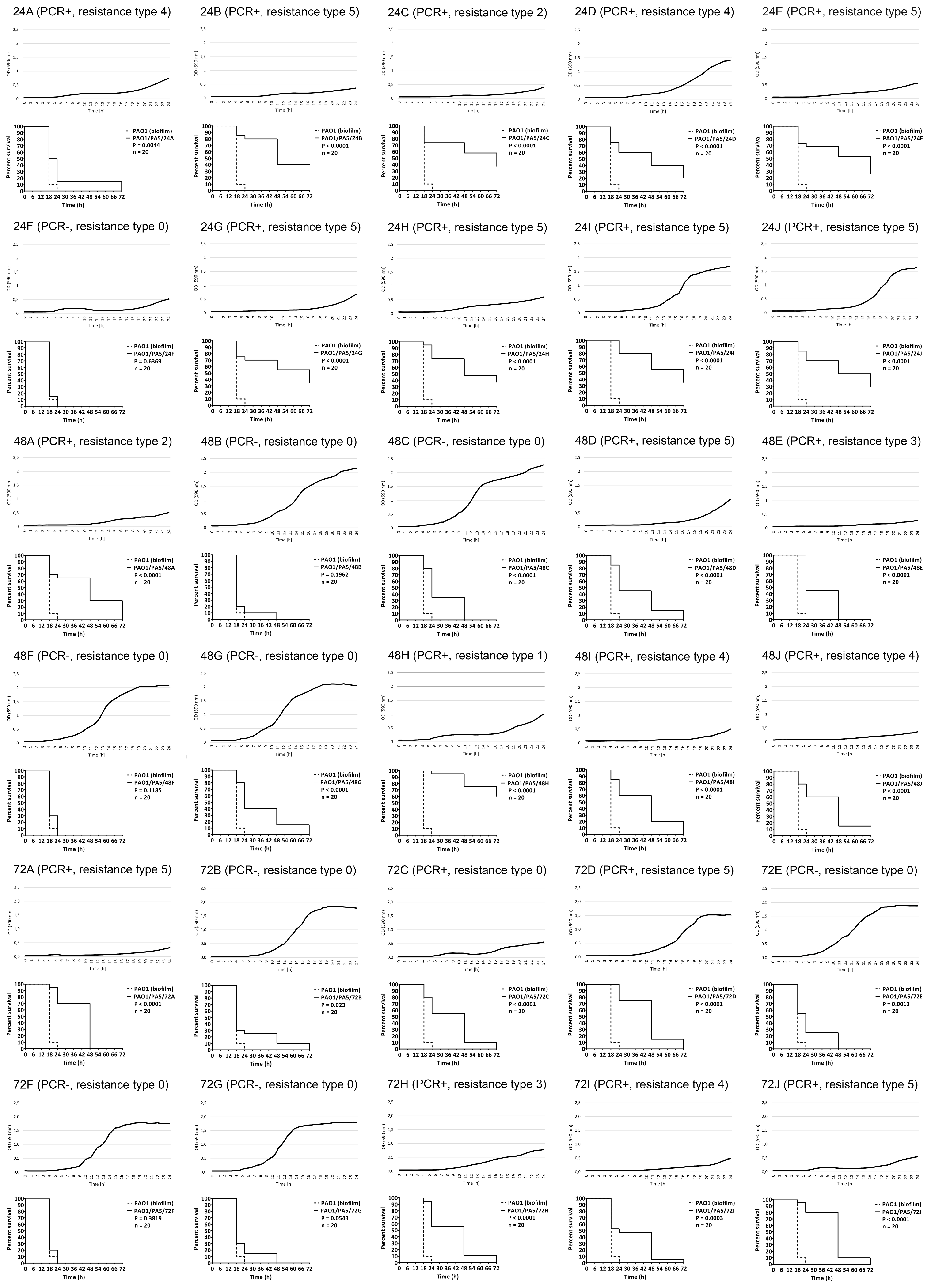

### Supplemental Fig. S3

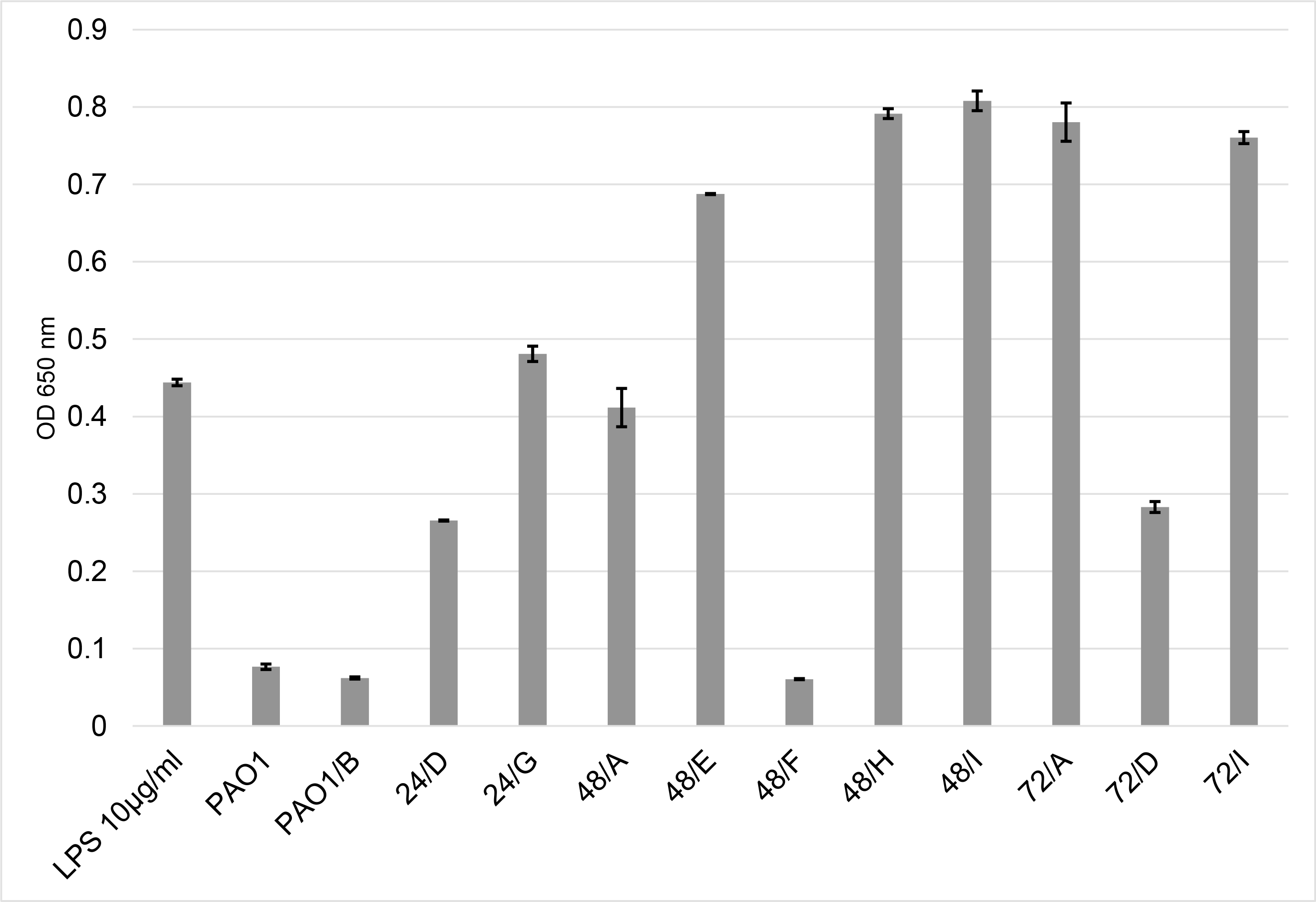
