## Supplemental Table S1 for "*Pseudomonas aeruginosa* PA5oct jumbo phage impacts planktonic and biofilm population and reduces its host virulence"

**Table S1**. Main features of *Pseudomonas* phages used in the study

| **Name** | **Taxonomy** | **Recognized bacterial receptor** |
| --- | --- | --- |
| PA5oct | myovirus, giant** | LPS/IV type pili |
| 14/1 | Pbunavirus, myovirus* | LPS |
| phiKZ | Phikzvirus, myovirus, giant* | IV type pili |
| LUZ19 | Phikmvvirus, podovirus* | IV type pili |
| LKD16 | Phikmvvirus, podovirus* | IV type pili |
| KMV | Phikmvvirus, podovirus* | IV type pili |

*Laboratory of Gene Technology, KU Leuven, Leuven, Belgium

**Department of Pathogen Biology and Immunology, Institute of Genetics and Microbiology, University of Wroclaw, Wroclaw, Poland
