## Supplemental Table S2 for "*Pseudomonas aeruginosa* PA5oct jumbo phage impacts planktonic and biofilm population and reduces its host virulence"

**Table S2**. Phage activity comparison of six virulent phages on *P. aeruginosa* strains from the University hospital of Leuven, Leuven, Belgium collection.

|  | **PA5oct** | **14/1** | **PhiKZ** | **LUZ19** | **LKD16** | **PhiKMV** |
| --- | --- | --- | --- | --- | --- | --- |
| **PAO1** | **+** | **+** | **+** | **+** | **+** | **+** |
| **PaLo1** |  |  |  |  |  |  |
| **PaLo2** |  |  |  |  |  |  |
| **PaLo3** | **+** |  |  |  |  |  |
| **PaLo4** | **+** |  |  |  |  |  |
| **PaLo5** | **+** |  | **+** |  |  | **+** |
| **PaLo6** | **+** |  | **+** |  |  | **+** |
| **PaLo7** |  | **+** |  |  |  | **+** |
| **PaLo8** |  | **+** |  | **+** | **+** |  |
| **PaLo9** |  | **+** |  | **+** |  |  |
| **PaLo10** | **+** |  |  |  |  | **+** |
| **PaLo11** | **+** |  |  |  |  |  |
| **PaLo12** | **+** |  |  |  |  |  |
| **PaLo13** |  |  | **+** |  |  |  |
| **PaLo14** | **+** |  |  |  |  | **+** |
| **PaLo15** | **+** | **+** | **+** |  |  |  |
| **PaLo16** |  |  |  |  |  |  |
| **PaLo17** |  | **+** |  |  |  |  |
| **PaLo18** | **nt** | **nt** | **nt** | **nt** | **nt** | **nt** |
| **PaLo19** |  |  |  |  |  |  |
| **PaLo20** |  |  |  |  |  |  |
| **PaLo21** | **+** |  | **+** | **+** | **+** | **+** |
| **PaLo22** | **+** |  |  |  |  |  |
| **PaLo23** |  |  |  |  |  |  |
| **PaLo24** |  |  |  |  |  |  |
| **PaLo25** |  |  |  |  |  |  |
| **PaLo26** |  | **+** |  |  |  |  |
| **PaLo27** |  |  |  |  |  | **+** |
| **PaLo28** |  |  |  |  |  |  |
| **PaLo29** |  |  |  |  |  |  |
| **PaLo30** |  |  |  |  |  |  |
| **PaLo31** |  |  |  | **+** | **+** | **+** |
| **PaLo32** |  |  |  |  |  |  |
| **PaLo33** |  |  |  |  |  |  |
| **PaLo34** |  |  |  |  |  |  |
| **PaLo35** |  |  |  |  |  |  |
| **PaLo36** | **+** | **+** | **+** | **+** | **+** | **+** |
| **PaLo37** |  |  |  |  |  |  |
| **PaLo38** |  |  |  |  |  |  |
| **PaLo39** | **+** |  | **+** | **+** | **+** | **+** |
| **PaLo40** |  | **+** |  |  |  |  |
| **PaLo41** |  |  |  |  |  |  |
| **PaLo42** | **+** |  |  | **+** | **+** | **+** |
| **PaLo43** | **+** |  |  | **+** | **+** | **+** |
| **PaLo44** | **+** |  | **+** | **+** | **+** | **+** |
| **PaLo45** |  |  |  |  |  |  |
| **PaLo46** | **+** |  |  |  |  |  |
| **PaLo47** | **+** | **+** |  |  |  |  |
| **# of sensitive strains** | **19** | **10** | **9** | **10** | **9** | **14** |
| **% of sensitive strains** | **40%** | **21%** | **19%** | **21%** | **19%** | **30%** |

nt – not tested
